# Intelligent SEEQPI enables reconstruction of inhibitory neuronal nuclei and dry-mass mapping deep in the living mouse brain

**DOI:** 10.64898/2026.10.25.761694

**Authors:** Jiaqi Yao, Jinhua Lyu, Naichen Shi, Xi Chen

## Abstract

Multiphoton fluorescence microscopy enables deep brain imaging with subcellular resolution and molecular specificity, but limited photon budgets constrained by tissue heating and nonlinear damage restrict temporal resolution of structural imaging in deeper regions. Moreover, imaging multiple molecularly defined cell populations requires additional fluorescent labels, while spectral bandwidth and crosstalk constrain multiplexed detection. Here we develop an intelligent imaging framework that uses label-free Scattering-Enabled Epi-Quantitative Phase Imaging (SEEQPI) to infer molecularly specific information deep *in vivo*. Using paired SEEQPI and three-photon fluorescence images, we developed a Feature-wise Linear Modulation U-Net with generative adversarial learning to reconstruct inhibitory-neuron nuclei. The network accurately predicts nuclear distributions and maintains performance when training and validation volumes are separated axially by up to 6 μm, enabling sparse fluorescence supervision. Cross-animal training further improves predictions in sparsely sampled target animals. Inferred nuclear maps enable dry-mass density estimation, demonstrating molecularly specific, label-free quantitative imaging deep within the living brain.

## Introduction

*In vivo* deep-tissue samples comprise complex biological structures with diverse morphologies that reflect their underlying biological functions^1,2^. Resolving these subcellular structures deep within tissue remains challenging^3,4^. Refractive-index heterogeneity causes strong light scattering and wavefront aberrations, which progressively distort and attenuate the optical field with increasing imaging depth^5–9^. Multiphoton microscopy is a powerful approach for high-resolution deep-tissue imaging by exploiting the reduced scattering of longer-wavelength excitation light and the nonlinear confinement of fluorescence to the focal volume, thereby enabling optical sectioning and high signal-to-background ratios at greater depths^10,11^. First reported in 1990, the two-photon laser scanning fluorescence microscope has been instrumental in demonstrating calcium dynamics in the dendrites of rat cortical neurons^12,13^. Building on this foundation, advances in ultrafast lasers have propelled three-photon microscopy to imaging depths beyond the reach of two-photon techniques, enabling successful *in vivo* imaging of deeper regions^14–16^. However, the low three-photon excitation probability necessitates high peak photon flux to generate sufficient fluorescence signal^17,18^. The resulting limited photon budget, further constrained by tissue heating and nonlinear photodamage, necessitates longer pixel dwell times or frame averaging, thereby limiting the temporal resolution of deep-tissue structural imaging^19–21^. Moreover, like other fluorescence-based approaches, three-photon microscopy relies on exogenous or genetically encoded fluorescent labels, which may perturb biological processes and are susceptible to photobleaching and phototoxicity during repeated or prolonged imaging^22,23^.

In contrast, linear-scattering-based label-free methods, which derive contrast from refractive index differences, require minimal illumination power^24–26^. This substantially reduces phototoxicity, eliminates photobleaching, and affords a higher photon budget for high temporal resolution, preserving the intact tissue microenvironment for long-term cellular investigation. Reflectance confocal microscopy uses a pinhole to reject out-of-focus light and improve axial sectioning, enabling deep *in vivo* mouse brain imaging at depths up to 1.3 mm^27^. However, lacking interferometric contrast enhancement, its intensity-based signal predominantly reflects large refractive index mismatches and struggles to resolve finer subcellular detail. Optical coherence microscopy (OCM) instead uses low-coherence interferometry, deriving contrast from interference between backscattered and reference light to achieve optical sectioning^28^. In deep-tissue imaging, however, dispersion in the reference and sample arms must be carefully matched and varies with depth, making the interferometric signal susceptible to noise and instability, and rendering phase retrieval particularly difficult. Therefore, although OCM can visualize structures such as vascular networks and myelinated axons, the cellular morphological contrast it provides remains limited.

To address these limitations, we developed Scattering-Enabled Epi-Quantitative Phase Imaging (SEEQPI), a label-free approach for live deep-tissue imaging with millimeter-scale imaging depth, subcellular resolution, and nanoscale phase sensitivity^29^. SEEQPI combines common-path interferometry to minimize dispersion mismatch with scattering-enabled phase retrieval to suppress incoherent background, enabling robust quantitative phase imaging deep within tissue. Its phase contrast resolves nuclei, somata, and neurites across diverse brain cell types and enables quantitative dry mass mapping^30–32^. However, SEEQPI lacks molecular specificity. To address this gap, we integrated SEEQPI with three-photon fluorescence and third-harmonic generation (THG) microscopy to acquire co-registered phase, molecular distributions, and structural interfaces. Nevertheless, fluorescence-based imaging remains limited in temporal resolution and molecular multiplexing.

Recent advances in deep learning offer a way around these limitations: supervised models can learn mappings from label-free images to fluorescence signals, inferring molecular specificity. Convolutional neural networks trained on paired gradient light interference microscopy (GLIM) phase images and wide-field epifluorescence images have predicted DNA- and RNA-specific fluorescence in spheroids^33^. With co-registered laser-scanning GLIM and confocal images, deep learning has also predicted molecularly specific signals with confocal-level resolution in organoids up to ~300 µm deep^24^. Similar strategies have inferred fluorescence from brightfield images in on-chip human induced pluripotent stem cell cultures^34^ and predicted nuclear and membrane fluorescence from transmitted-light images in live mouse intestinal organoids^35^.

Deep learning has also been combined with sample-expansion stimulated Raman scattering for label-free, super-resolution volumetric identification of multiple cellular structures in mouse brain tissue^36^. More recently, approaches have moved beyond U-Net-based regression toward generative and task-specific frameworks for complex 3D data, including generative adversarial network (GAN)-based reconstruction of tissue microstructures and neural-network-based probabilistic analysis of densely packed organoids^37,38^. Nevertheless, inferring molecularly specific fluorescence from label-free images of intact, deeply embedded living mammalian brain tissue remains largely unexplored, because extensive cellular and structural heterogeneity makes reliable inference difficult.

In this work, we present an intelligent SEEQPI framework that predicts three-photon fluorescence from inhibitory-neuron nuclei in the deep living mouse brain. We trained a two-stage model on co-registered SEEQPI and three-photon fluorescence images acquired *in vivo* at depths up to 1.1 mm. Supervised regression first established nuclear localization with a Feature-wise Linear Modulation U-Net (FiLM U-Net), and conditional GAN fine-tuning with PatchGAN then sharpened nuclear contours. The FiLM U-Net takes three adjacent SEEQPI images and the central plane’s z-coordinate as inputs to improve axial sectioning, while PatchGAN evaluates local statistical realism to enhance lateral contrast. On randomly partitioned images from multiple mice, the model achieved high sensitivity and precision, establishing translational feasibility. Within a single mouse, training on planes spaced up to 6 μm apart enabled prediction of nuclear distributions at intermediate depths, and cross-animal learning further improved sensitivity. This supports a hybrid strategy that acquires three-photon images only at sparse planes and predicts the planes between them, improving deep-tissue acquisition efficiency and enabling faster time-lapse imaging. Combined with deep-learning inference, intelligent SEEQPI enables high-resolution quantification of dry mass associated with specific molecular structures deep within living tissue.

## Results

### Paired SEEQPI and three-photon image acquisition

We constructed a multimodal imaging system to acquire coregistered SEEQPI, three-photon fluorescence, and THG images^29^ (Fig. 1a, see Methods for details). SEEQPI uses common-path phase-shifting interferometry implemented in a confocal differential interference contrast (CDIC) configuration. Each SEEQPI image is reconstructed from four CDIC frames acquired at 90° phase increments using the scattering-enabled self-referenced algorithm^29^ (Fig. 1b). Figure 1c compares the three imaging modalities at different imaging depths in a living mouse brain. SEEQPI contrast arises from refractive index variations, enabling visualization of diverse cellular components. In contrast, THG highlights structural interfaces, while three-photon fluorescence visualizes inhibitory neuronal nuclei labeled through viral injections. However, labeling efficiency depends on factors including injection site, viral concentration, and the animal’s physiological condition; thus, some inhibitory neuronal nuclei may not be labeled or detected. In the following analysis, we refer to the three-photon fluorescence images as “ground truth” for brevity, although they represent a subset of the inhibitory neuronal nuclei.

**Fig. 1.**
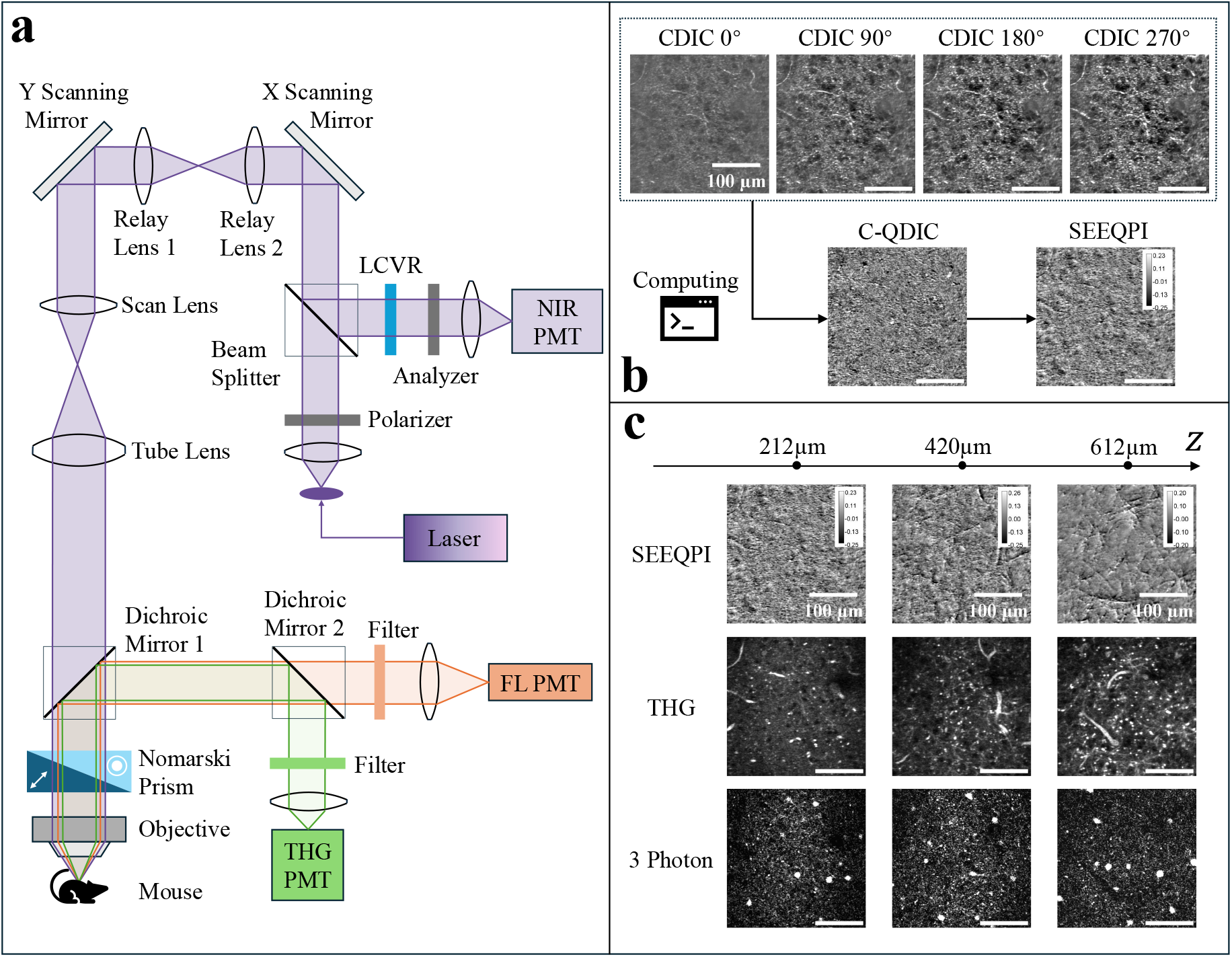
Workflow for simultaneous acquisition of coregistered SEEQPI, THG, and three-photon fluorescence images. **a**, The multimodal imaging system uses a common laser source for all three modalities. The SEEQPI optical path incorporates an input polarizer, a Nomarski prism, an analyzer oriented orthogonally to the input polarizer, and a pinhole, forming a confocal differential interference contrast (CDIC) configuration. Backscattered light from the sample passes through a liquid crystal variable retarder (LCVR) and the analyzer before detection by a near-infrared photomultiplier tube (PMT). Three-photon fluorescence and THG signals are separated from the excitation light and from each other using two dichroic mirrors and spectral filters, then detected by dedicated PMTs. **b**, Four CDIC images are acquired by sequentially adjusting the LCVR voltage to introduce phase shifts spaced 90° apart. These images are processed to obtain a confocal quantitative differential interference contrast (C-QDIC) image, from which the SEEQPI phase map is reconstructed using the scattering-enabled, self-referenced algorithm. **c**, Coregistered SEEQPI, THG, and three-photon fluorescence images are compared at different imaging depths.

Volumetric z-stacks were acquired from four mice to depths of ~1100 μm (imaging parameters in Fig. 2a). Inhibitory-neuron nuclei masks were generated by thresholding the three-photon images (preprocessing details in Methods). The z-stacks were partitioned into training, validation, and test sets using three schemes. In Scheme 1 (Fig. 2b), slices from all four mice were pooled and randomly assigned to the three sets, establishing a baseline for overall translation performance. In Scheme 2, we used the z-stack from a single mouse, with images sampled every Δ*z* μm forming the training set and the remainder split between validation and testing (Fig. 2c, Δ*z* is 4 μm), evaluating reconstruction of intermediate slices under sparse three-photon sampling. Scheme 3 (Fig. 2d) retained Scheme 2 split for the target mouse and added training images from the other three mice sampled at the same spacing, evaluating cross-mouse generalization.

**Fig. 2.**
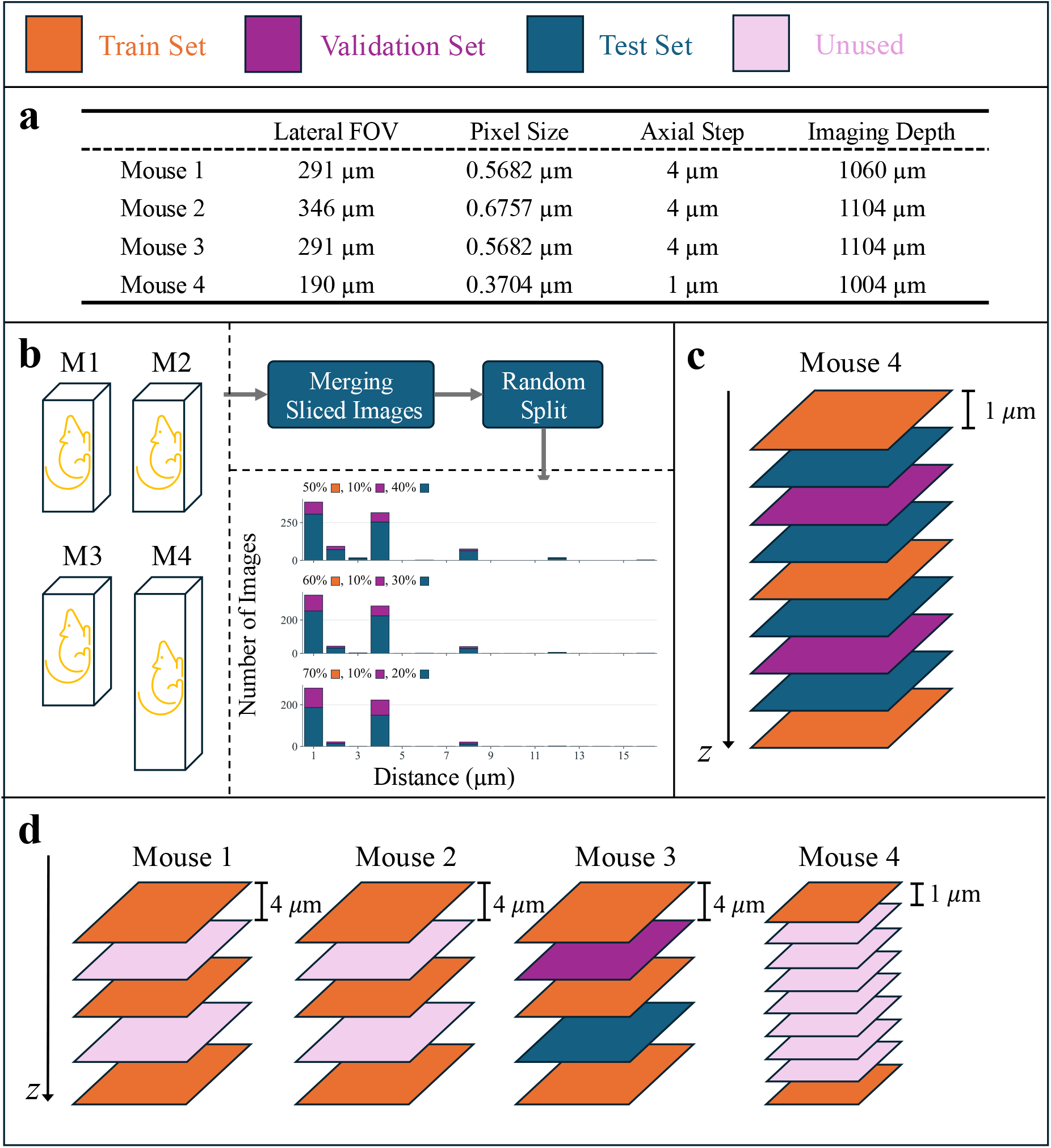
Acquisition parameters for z-stacks from four mice and data partitioning strategies for training, validation, and testing. **a**, Z-stacks from Mice 1–3 were acquired over a larger field of view (FOV) with a 4-μm axial step size. Mouse 4 was imaged over a smaller FOV and with 1-μm axial step size. **b**, Data partitioning Scheme 1: images from all four mice were pooled and randomly divided into training, validation, and test sets according to the specified ratios. **c**, Data partitioning Scheme 2: images from a single mouse were sampled at a fixed axial interval Δ*z* for training, and the remaining images were randomly divided into validation and test sets. The example shows a sampling interval Δ*z* of 4 μm. **d**, Data partitioning Scheme 3: the training set from Scheme 2 was expanded by adding images sampled at the same axial interval Δ*z* from the other three mice. The remaining images from these additional mice were unused, while the validation and test sets were drawn exclusively from the target mouse. The example shows a sampling interval Δ*z* of 8 μm, with Mouse 3 as the target.

Deep neural networks were trained for image-to-image translation from SEEQPI to three-photon images. At each depth *z*, the network takes three adjacent SEEQPI images and the central plane’s *z*-coordinate to predict the corresponding three-photon image, which is thresholded to yield a binary nuclear mask (Fig. 3a). SEEQPI images show dense structural detail continuously to 1,100 μm, whereas three-photon images show sparse nuclear signals mainly from the surface to ~800 μm, above the white matter. The network comprises a FiLM U-Net generator and a PatchGAN discriminator (Fig. 3b). Training proceeds in two stages: the FiLM U-Net is first trained by supervised learning^39,40^, then fine-tuned within a conditional GAN, where PatchGAN evaluates the local statistical realism of generated images^41^. The resulting models are hereafter called FiLM U-Net and FiLM U-Net + GAN, and both are retained for inference (details in Methods).

**Fig. 3.**
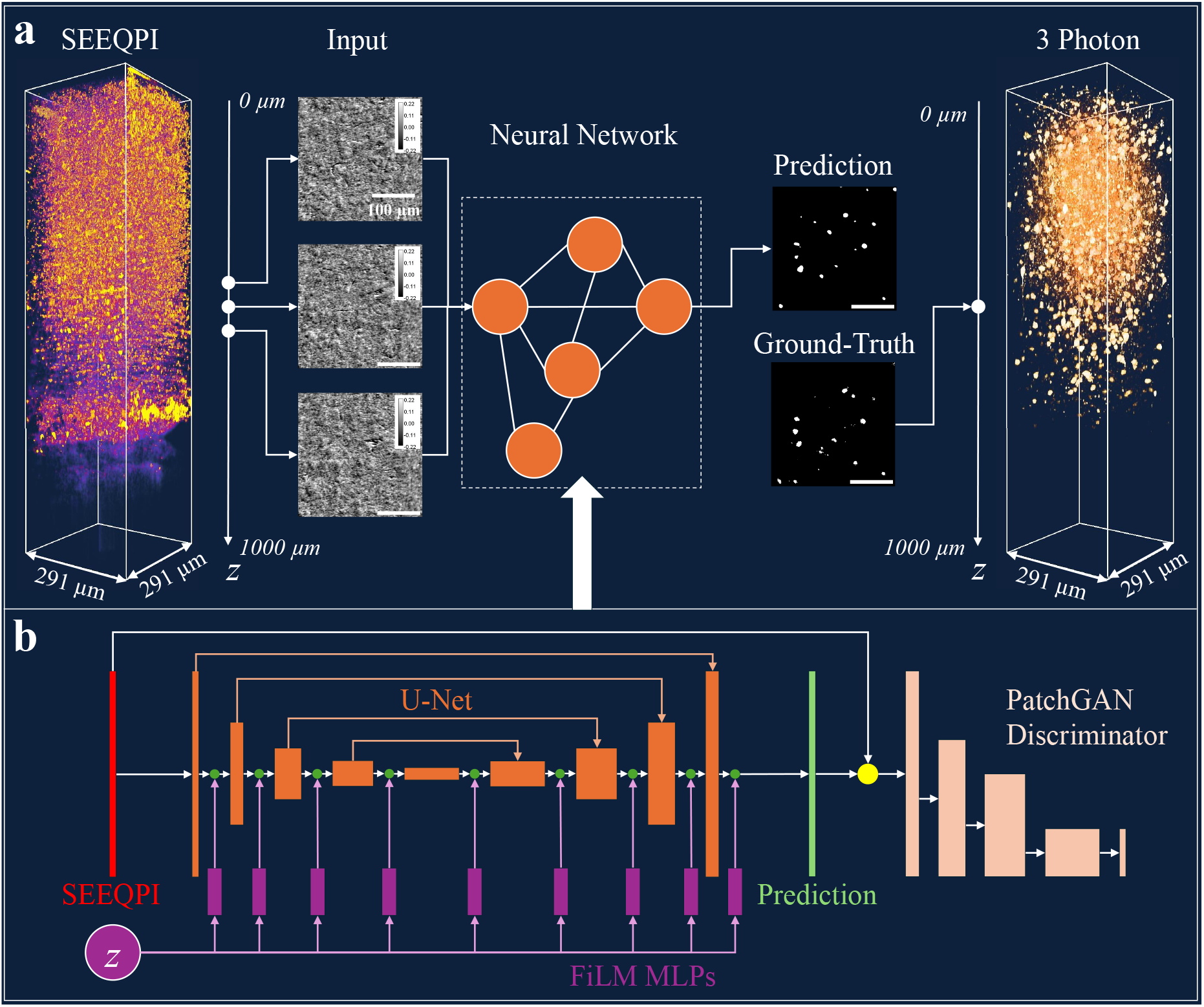
The framework of image translation. **a**, Schematic diagram of the pipeline from z-stack SEEQPI images to 3-photon masks. For each position within the z-stack, three adjacent SEEQPI images and their corresponding z-axis coordinates are fed into the neural network, which then outputs the predicted 3-photon mask. **b**, Detailed architecture of the neural network. The SEEQPI images are fed into the U-Net as a 3-channel input. Simultaneously, the z-axis coordinates are processed by a multilayer perceptron (MLP) and encoded into the outputs of each convolutional block within the U-Net, forming the complete FiLM U-Net architecture. Subsequently, the U-Net output is passed to a PatchGAN discriminator to evaluate its authenticity, providing a feedback signal to further refine the FiLM U-Net.

### Three-photon reconstruction of inhibitory neuron

We first pooled the sliced images from four mice’s z-stacks, and then proportionally partitioned them into the training, validation, and test sets. Three distinct allocation ratios for these three sets were employed: 70%/10%/20%, 60%/10%/30%, and 50%/10%/40%. An input layer dropout strategy was implemented to improve the networks’ generalization, as described in the Results Section. The dropout probability, defined as the probability of replacing an adjacent input channel with zeros, was set to 0.05.

In Fig. 4a, we visualized the 2D predicted nuclei masks across different mice under the 70%/10%/20% configuration. The mask removes background noise while preserving larger connected clusters in raw three-photon images. Comparison with the ground-truth masks shows that both models closely reproduce the spatial distribution of neuronal nuclei, supporting their predictive accuracy. However, FiLM U-Net identifies more false nuclear regions than FiLM U-Net + GAN, particularly in Mice 1-3. Conversely, as highlighted by the red box in Mouse 4, FiLM U-Net + GAN recovers additional true nuclear signals missed by FiLM U-Net. This confirms that GAN fine-tuning improves the prediction accuracy.

**Fig. 4.**
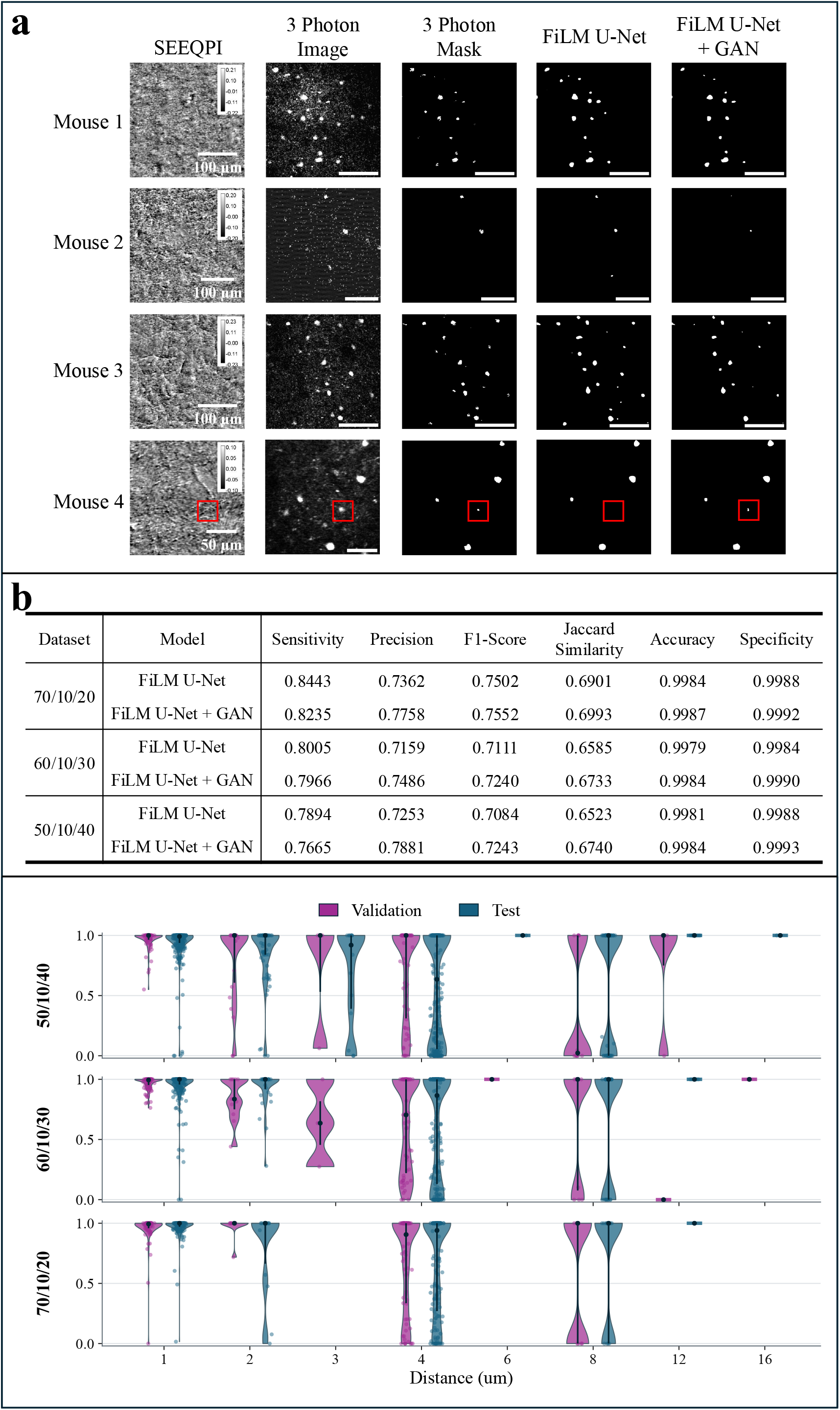
Prediction results in Scheme 1. **a**, Representative 2D image slices from the test set obtained under the 70%/10%/20% configuration. From left to right, the five columns display the SEEQPI image, the corresponding ground-truth three-photon fluorescence image, the binary mask derived from three-photon images, the predicted three-photon mask from FiLM U-Net, and the predicted three-photon mask from FiLM U-Net + GAN, all corresponding to the same FOV and z-axis coordinate. From top to bottom, the four rows display representative images selected from Mouse 1 through Mouse 4, respectively, illustrating models’ prediction results across different mice. The corresponding z-axis locations for these four selected images within their respective volumes are 200 µm, 464 µm, 376 µm, and 414 µm. **b**, Quantitative evaluation of model predictions on the test set. Performance is compared between the FiLM U-Net and GAN models across the three configurations using accuracy, sensitivity, specificity, precision, F1-score, and Jaccard similarity. **c**, Violin plots showing the relationship between prediction sensitivity and distance across the three configurations. Distance is defined as the physical distance between the evaluated image and its nearest counterpart in the training set.

In Fig. 4b, we quantitatively evaluated the performance of both FiLM U-Net and FiLM U-Net + GAN models across the three configurations, with the definitions of the used metrics in the Methods Section. Because inhibitory neurons visible in three-photon images represent only a subset of the true population, some metrics are less informative. Sensitivity was prioritized because neurons identified by three-photon imaging served as true positive targets for evaluating the model’s ability to detect inhibitory neuronal nuclei. In contrast, predicted inhibitory neuronal nuclei not detected in the three-photon fluorescence images may still be present in the tissue, limiting the reliability of precision estimates. Nevertheless, precision is retained as a reference metric for assessing whether predictions preserve the statistical characteristics of three-photon images. Across all three training configurations, the models achieved sensitivities of 0.75–0.85, indicating that they detected the majority of inhibitory neuronal nuclei identified by three-photon fluorescence imaging. Missed boundary pixels likely prevented sensitivity from reaching 1. When comparing the three configurations, we observed that sensitivity decreased as the proportion of the training set was reduced. This trend may be explained by poorer generalization due to insufficient coverage of the data distribution in the training set. Within the same configuration, FiLM U-Net has a higher sensitivity but lower precision than FiLM U-Net + GAN, indicating that the latter yields more conservative predictions regarding neuronal locations. The F1-score and Jaccard similarity, which jointly reflect sensitivity and precision, are higher for the GAN model, suggesting closer overall agreement with the ground truth. Due to the sparse nature of nuclei spatial distribution, both accuracy and specificity maintain near-perfect values across three configurations.

To identify factors affecting single-image sensitivity, Fig. 4c relates sensitivity to the physical distance between each evaluated image and its nearest training image across the three configurations. Within each configuration, predicted sensitivity distributions shifted toward lower values with increasing distance in both the validation and test sets. The median exceeds 0.8 at a distance of 1, whereas many instances fall below 0.5 at a distance of 8. Across the three configurations, the overall sensitivity distribution improves with the proportion of the training set increases, as test images are closer to the training data. These results suggest that the depth difference between test and training images is an important factor influencing prediction performance.

### Three-photon reconstruction with sparse axial supervision

In the second scheme, within a single mouse z-stack, images sampled at constant intervals Δ*z* formed the training set, while the remainder were split into validation and test sets with the configuration shown in Fig. 2c. In Mouse 4, experiments were conducted at Δ*z* = [4,5,6,7,8] μm, with corresponding dropout probabilities of [0.05, 0.10, 0.12, 0.12, 0.12].

Figure 5a presents a comparison of the ground-truth and predicted two-dimensional images at different depths when Δ*z* = 4 μm. Both FiLM U-Net and FiLM U-Net + GAN accurately predicted nuclei present in the ground-truth nuclei mask while also identifying additional nuclear regions, particularly at greater imaging depths. Compared with FiLM U-Net, FiLM U-Net + GAN recovered small clusters of putative nuclear signals that were weak or absent in the three-photon masks and may represent inhibitory neurons not captured by three-photon imaging., as illustrated at a depth of 120 μm. In Fig. 5b, the predicted masks from the test set were converted into three-dimensional renderings. The ground-truth, results of FiLM U-Net, and FiLM U-Net + GAN exhibit highly consistent spatial distributions, demonstrating our models’ ability to reconstruct inhibitory neuronal nuclei distribution in brain tissue at different imaging depths.

**Fig. 5.**
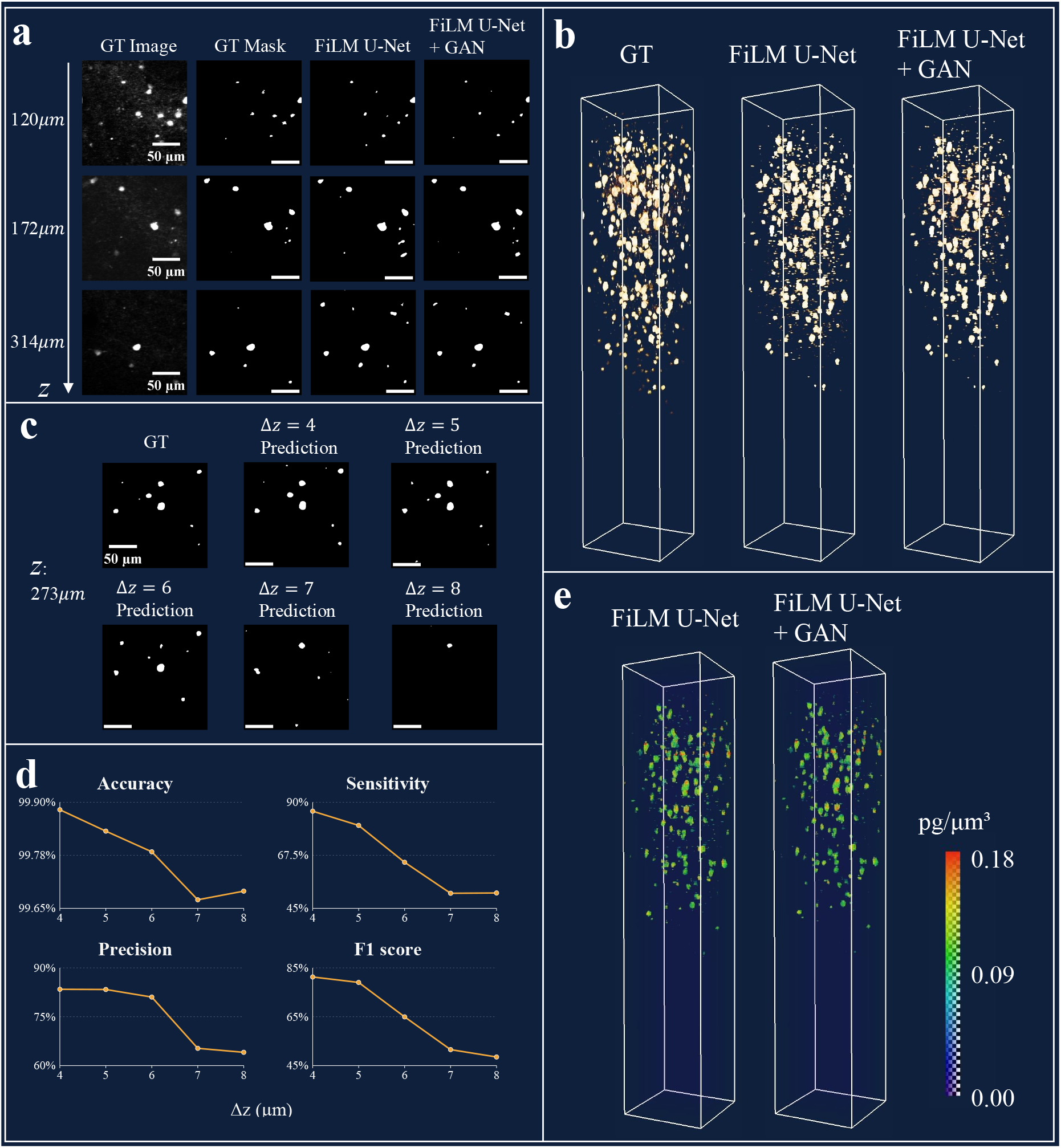
Nuclei mask prediction and dry mass mapping in Scheme 2. **a**, Comparison of ground-truth and model-predicted 2D images for Mouse 4. From left to right, the columns display the ground-truth image, the ground-truth mask, the predicted mask from the FiLM U-Net model, and the predicted mask from the fine-tuned GAN model. From top to bottom, the rows represent imaging depths of 120 µm, 170 µm, and 314 µm, respectively. **b**, Comparison of 3D-rendered 3-photon masks among the ground truth, the FiLM U-Net model, and the GAN model. Since all images in the z-stack were drawn from the test set, the 3D rendering appears relatively flattened, with the predicted signals concentrated at shallower depths. **c**, Comparison of the ground-truth 3-photon mask and the GAN model predictions at a depth of 273 µm as Δ*z* varies from 4 µm to 8 µm. **d**, Line graphs illustrating the variations in the model’s predictive accuracy, sensitivity, precision, and F1-score as Δ*z* increases from 4 to 8 µm. **e**, 3D renderings of the dry mass density distribution of inhibitory neurons, estimated by the FiLM U-Net and GAN models. The models predict the spatial locations of the neurons, while the dry mass density at these specific locations is calculated using the SEEQPI phase maps.

However, reconstruction sensitivity and precision are expected to decrease with Δ*z* since it depends on the distance to the nearest training image, as shown above. Hence, we sought to determine the maximum Δ*z* at which prediction performance remains acceptable. As shown in Fig. 5c, at the same depth, we compared the masks predicted by FiLM U-Net + GAN with the ground-truth for Δ*z* ranging from 4 to 8 μm. When Δ*z* was 4 or 5 μm, predictions closely matched the ground truth. At 6 μm, larger nuclei remained detectable, whereas smaller ones were often missed; performance deteriorated substantially at 7-8 μm. Figure 5d confirms this trend: although accuracy remained high, sensitivity and F1-score declined with Δ*z*, while precision remained above 0.75 only up to 6 μm. In conclusion, we selected 6 μm as the maximum acceptable training interval.

The SEEQPI images and predicted three-photon masks were further used to quantify the dry mass density of inhibitory neurons, with the method presented in the Methods Section. The predicted masks delineated nuclear locations and boundaries, while SEEQPI estimated dry-mass density through the linear relationship between phase delay and dry mass. Figure 5e presents 3D renderings of the dry mass density distributions of inhibitory neurons predicted by both neural networks. The estimated nuclear dry mass densities ranged from 0.09 to 0.18 pg/μm^3^, consistent with previously reported experimental measurements. This agreement supports the ability of our intelligent SEEQPI framework to accurately quantify the dry mass density of inhibitory neurons.

### Cross-mouse generalization with multi-mouse training

This scheme evaluates models’ cross-mouse generalization. Given the limited performance of single-mouse training at large Δ*z* in Scheme 2, we tested whether incorporating images from multiple mice could improve prediction accuracy. Using Mouse 3 as the target, we followed Scheme 2 with Δ*z* = 8 μm. For comparison, a single-stack model was trained only on the sparsely sampled Mouse 3 train set, whereas a multi-stack model additionally included images sampled at the same interval Δ*z* = 8 μm from other mice for training. Both models were evaluated on the same Mouse 3 test set. The input layer dropout probability is 0.15.

Figure 6a compares the ground-truth three-photon images with the corresponding predictions generated by the single-stack and multi-stack learning models. As shown, the spatial distribution of nuclei predicted by the single-stack learning model was poorly aligned with that observed in the ground-truth mask, which is consistent with our findings at Δ*z* = 8 μm in Scheme 2. In contrast, multiple-stack learning substantially increased the number of predicted nuclei, many of which corresponded to nuclei present in the ground-truth, resulting in a marked improvement in sensitivity. Moreover, the model produced substantially more signals than observed in the three-photon images, some of which may correspond to true inhibitory-neuron locations identified by the model but not captured by three-photon imaging.

**Fig. 6.**
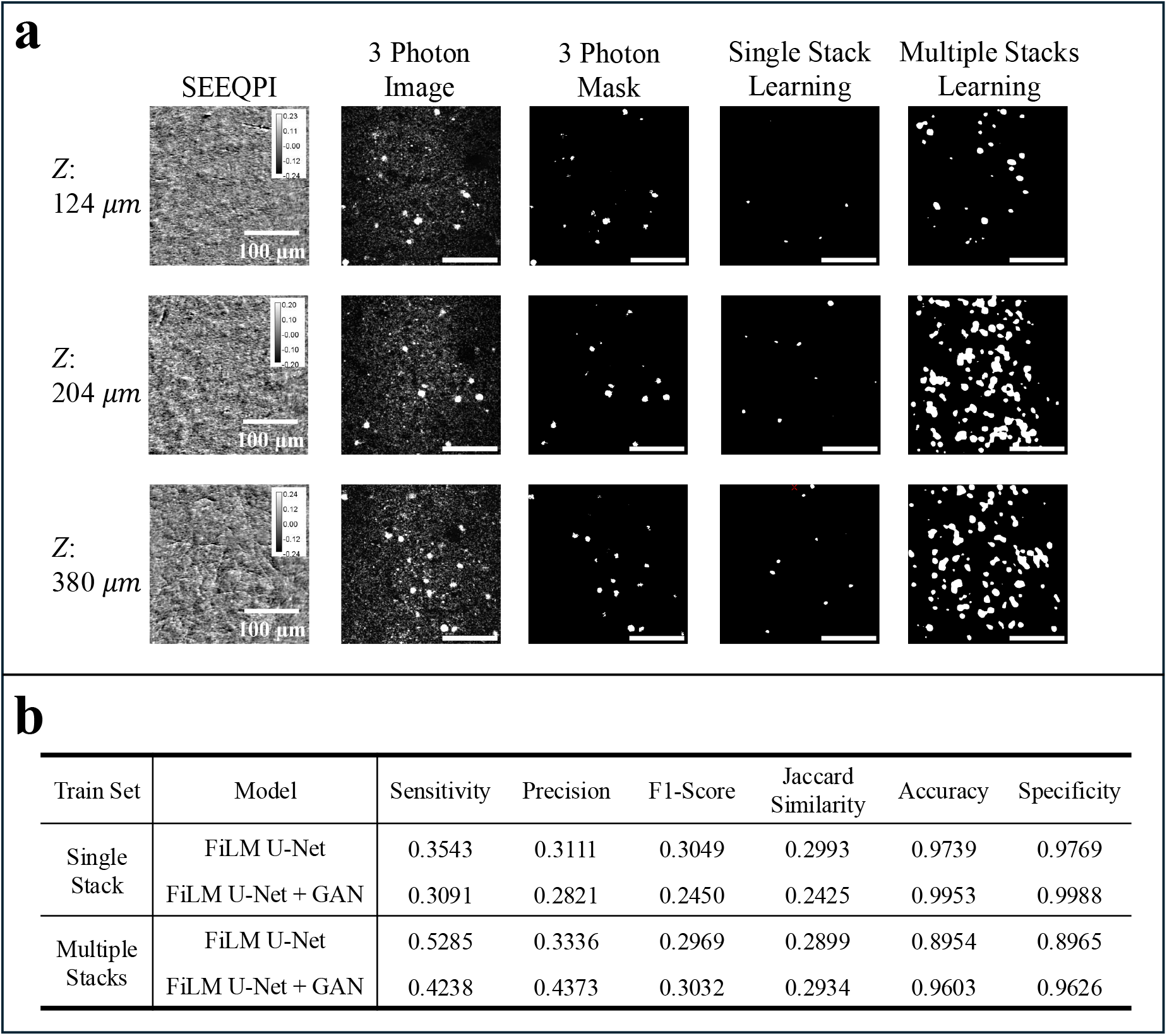
Prediction performance of the models on the test set in Scheme 3. **a**, Comparison of the SEEQPI, ground-truth, and predicted 2D images at various depths in Mouse 3. From left to right, the columns display the SEEQPI image, the ground-truth 3-photon image, the ground-truth mask, the predicted mask from the single-stack learning GAN model, and the predicted mask from the multiple-stack learning GAN model. From top to bottom, the rows present images at depths of 124 µm, 204 µm, and 380 µm, respectively. **b**, Statistical analysis of the prediction results under single-stack and multiple-stack learning. The following metrics are used: accuracy, sensitivity, specificity, precision, F1-score, and Jaccard similarity.

Figure 6b further quantified the performance of FiLM U-Net and FiLM U-Net + GAN under both single-stack and multi-stack learning with our defined metrics. Compared with the two single-stack models, multi-stack learning markedly improved sensitivity, particularly for the supervised FiLM U-Net. This improvement, however, was accompanied by substantial decreases in accuracy and specificity. GAN fine-tuning largely mitigated this overprediction, restoring accuracy and specificity to high levels, although sensitivity decreased slightly after fine-tuning. Interestingly, both multi-stack models achieved substantially higher precision than their single-stack counterparts, indicating that more of the additional predicted signals corresponded to true nuclei. Consistent with these improvements, multi-stack learning also yielded higher F1 scores and Jaccard similarity.

## Discussion

Deep-tissue fluorescence imaging provides the molecular specificity needed to identify particular cell populations and compartments in living tissue. At greater depths, however, weak signals often require prolonged averaging, which limits temporal resolution, or increased excitation power, which raises concerns about heating, photobleaching, and phototoxicity. Multiplexing is also constrained by the limited number of fluorescence channels and labeling strategies. SEEQPI offers a complementary approach. Its contrast arises from refractive-index variations within tissue, enabling label-free imaging of diverse cellular and subcellular structures and providing quantitative phase information related to morphology and dry mass. However, SEEQPI does not directly provide molecular specificity, so specific neuronal types or compartments are difficult to identify from SEEQPI images alone.

To bridge these capabilities, we developed a deep-learning framework that translates SEEQPI images into three-photon fluorescence-like images, predicting inhibitory-neuron nuclear locations deep within the living mouse brain. This is challenging because the brain contains heterogeneous cell populations with overlapping morphologies, while scattering and depth-dependent contrast changes further complicate discrimination. Nevertheless, the network learned to predict these locations from *in vivo* SEEQPI images, suggesting that SEEQPI encodes cell-type-associated information not readily apparent by visual inspection, possibly through phase cues related to morphology and dry mass. We also combined the predicted nuclear masks with the phase–dry-mass relationship to estimate the nuclear dry-mass density of inhibitory neurons. The resulting distributions agreed with experimental measurements, extending label-free imaging from structural identification to quantitative cellular analysis.

We designed three schemes with distinct prediction objectives to evaluate feasibility and practical utility. In Scheme 1, pooled z-stack images from four mice were randomly divided into training, validation, and test sets. The high sensitivity and precision demonstrated the feasibility of SEEQPI-to-three-photon translation, although performance depended on the distance between target and training images. Scheme 2 predicted intermediate planes within a single-mouse z-stack from sparsely sampled training images; 6 μm was the largest tested interval that maintained satisfactory performance. These findings support a hybrid acquisition strategy for deep-tissue volumetric imaging. Where three-photon acquisition requires averaging over tens of seconds, SEEQPI is substantially faster because its photon budget is orders of magnitude higher. Three-photon images could therefore be acquired at selected axial positions, with SEEQPI images at intermediate positions used for network prediction. This could reduce fluorescence acquisition time, enable denser sampling within shorter windows, and reduce temporal variation across axial planes in time-lapse imaging. The 6 μm limit, however, constrains the potential efficiency gain.

Scheme 3 showed that data from other mice improved prediction sensitivity and precision in a target mouse when only sparse target-specific training images were available, suggesting that some learned features transfer across animals despite the heterogeneity of living brain tissue. Generalization remained limited, however. Cellular distributions, tissue architecture, and contrast vary between animals, so features associated with inhibitory neurons in training mice may not reliably distinguish them in an unseen mouse. Indeed, a model trained exclusively on other mice could not accurately localize inhibitory-neuron nuclei in a new animal. Cross-mouse training can therefore supplement sparse target-specific supervision but not yet replace it. Limited generalization is the main challenge across all three schemes, and the framework cannot yet eliminate the need for fluorescence labeling or three-photon acquisition. These limitations nonetheless point toward reducing dependence on fluorescence rather than replacing one modality with another.

A further consideration is that three-photon fluorescence does not provide a complete ground truth for the inhibitory-neuron population, because identification depends on the labeling strategy and on reporter expression and detectability. The network is therefore trained to reproduce the detectable signal, not necessarily every inhibitory-neuron nucleus. This also creates an opportunity: because SEEQPI provides structural information distinct from fluorescence intensity, a network might identify inhibitory neurons beyond the subset visible in the training data. The present results do not demonstrate this, and it requires independent validation. Correlative *ex vivo* immunostaining could help assess completeness and specificity, but matching individual nuclei between *in vivo* imaging and fixed, sectioned tissue is difficult, and processing could alter morphology and registration. In future work, we plan to use genetically modified mice with more complete and specific labeling to provide a more reliable reference.

Several directions may advance this approach. First, transformer- and diffusion-based architectures may better capture spatial relationships in SEEQPI images, although they should be evaluated against appropriate baselines. Second, the weighted mean-squared-error loss emphasizes neuron-positive regions but does not distinguish inhibitory neurons from other cell types, so larger datasets with cell-type annotations and multiclass segmentation could improve specificity. Third, more animals may help the network learn features that generalize across individual differences. Finally, more comprehensive genetic labeling combined with correlative in vivo and ex vivo imaging could test whether SEEQPI-trained networks identify cells beyond those detected by fluorescence. Together, these developments may allow label-free quantitative imaging and deep learning to complement fluorescence microscopy, reduce prolonged or repeated fluorescence acquisition, and improve access to cellular information in living tissue.

## Methods

### Multimodal Imaging System Setup

We constructed a multimodal system that consists of three photon fluorescence, THG, and SEEQPI. The system employs a femtosecond laser with a center wavelength of 1650 nm and a pulse duration of approximately 70 fs. The beam then passes a dispersion compensator and a pair of single-axis X and Y galvanometric mirrors conjugated by relay lenses, which control the focal point on the sample. The scan lens and tube lens pair expands the beam diameter to 13 mm. After the beam is focused onto the sample by an objective, the generated SEEQPI, three-photon, and THG signals are back-collected by the same objective. Details of the optical components are provided in the SEEQPI publication^29^.

In the SEEQPI pathway, the polarizer produces linearly polarized light, the Nomarski prism splits it into two orthogonally polarized beams probing slightly displaced sample positions, and the analyzer projects the returning beams onto a common polarization state to enable interference. As a linear system, the scattered light maintains the same wavelength as the incident light. The LCVR introduces a controlled phase shift to one of the linearly polarized components relative to the other. The image is captured by the NIR PMT. Simultaneously, as the laser beam illuminates the sample, it generates corresponding three-photon and THG emissions. As nonlinear processes, these emissions possess different wavelengths from the excitation light: 550 nm for THG, approximately 600 nm for three-photon fluorescence labeled with mRuby2, and approximately 647 nm for 7-AAD. Consequently, these signals can be effectively separated by two dichroic mirrors and their respective filters, and ultimately detected by their designated PMTs.

Deep tissue samples were obtained from mice that underwent craniotomy surgery. Subsequently, AAV1-mDlx-NLS-mRuby2 was injected to label inhibitory neurons within the region of interest. Detailed surgical procedures are shown in the SEEQPI publication^29^. The acquired SEEQPI, three-photon, and THG images had a spatial resolution of 512 × 512 pixels. The acquisition frame rate for SEEQPI was 0.27 Hz. Owing to frame averaging, the frame rates for THG and three-photon imaging were 0.27 Hz for superficial layers (< 500 μm) and 0.07 Hz for deeper layers (> 500 μm).

### Image normalization and segmentation

Both the SEEQPI and three-photon image stacks underwent per-stack normalization. The THG images were left unprocessed, as they were not utilized in the subsequent neural network training. Per-stack normalization was selected over per-slice normalization to avoid amplifying noise in deep regions. Z-score normalization was applied to SEEQPI images to prevent high-intensity outliers from skewing the overall signal distribution. Conversely, the three-photon images were normalized to the [0,1] range as 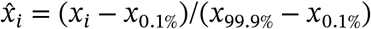, where *x*_99.9%_ and *x*_0.1%_ are the 99.9% and 0.1% quantile of pixel values. Pixel values falling outside the [0, 1] interval were clipped to 0 or 1. A global threshold of τ = 0.2 was used to generate the binary mask of 3-photon images. The resulting mask was further refined by removing 26-connected neighbors with a volume smaller than 300 voxels to eliminate background noise.

### Neural network architecture

Finally, a threshold τ^′^ = 0.2 is applied to generate a predicted binary mask, where positive pixels indicate the presence of a cell nucleus.

We employ the Pix2Pix conditional generation architecture within a generative adversarial network framework. In this setup, a FiLM U-Net **g**_**θ**_ serves as the generator, while a standard PatchGAN discriminator **d**_**ϕ**_ is employed to penalize structures at the scale of local image patches. The FiLM U-Net consists of a standard U-Net architecture and nine feature-wise linear modulation (FiIM) blocks. The U-Net utilizes a kernel size of 3, a stride of 2, and a padding of 1. Each FiLM layer employs a hidden layer with 128 neurons to generate scale and shift vectors, making an affine transformation to the intermediate feature map of U-Net. Using a sigmoid activation function, the predicted three-photon image **g**_**θ**_(**x**, *z*) is synthesized. Furthermore, the PatchGAN discriminator evaluates channel-concatenated pairs of the input SEEQPI images and either generated or ground-truth three-photon images, producing a patch-level realism map **d**_**ϕ**_(**x, y**).

The training process is divided into two stages. The first stage is pure FiLM U-Net supervised training. The network learns the image translation relationship from the paired ground-truth SEEQPI **x** and 3-photon images **y** with the weighted MSE loss function, which is defined as

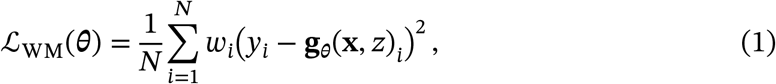

where *y*_*i*_ and **g**_**θ**_(**x**, *z*)_*i*_ are the ground-truth and predicted pixel values in the 3-photon image. *w*_*i*_ denotes the pixel-wise weight assigned to each individual pixel, representing its relative importance in updating the neural network parameters during training. In our training, *w*_*i*_ is a binary variable defined as

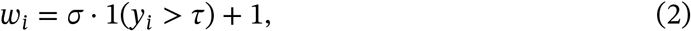

where τ is the threshold for generating the ground-truth 3-photon mask. σ controls the difference between two binary values, set as 100 in our experiments. The rationale for introducing spatial weights into the MSE loss stems from the highly sparse nature of real three-photon signals. In the first stage, the model was trained using the Adam optimizer with a learning rate 5 × 10^−4^, *β*_1_ = 0.9, *β*_2_ = 0.999 and an L2 regularization coefficient of 5 × 10^−9^.

In the second stage, the FiLM U-Net and the PatchGAN discriminator were alternately optimized within an adversarial framework. First, the discriminator’s ability to distinguish between synthesized and ground-truth three-photon images was improved with the following binary cross-entropy adversarial loss function

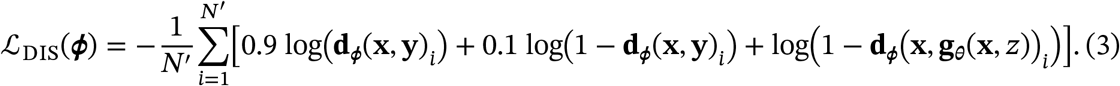

The target label for ground-truth image pairs is set as 0.9 to prevent overconfidence and improve stability. The discriminator is then optimized with an Adam optimizer with a learning rate 1 × 10^−5^, *β*_1_ = 0.5, *β*_2_ = 0.999. Second, we drove the FiLM U-Net to further refine the local physical details of the predictions with the composite loss function

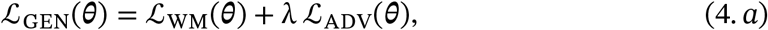

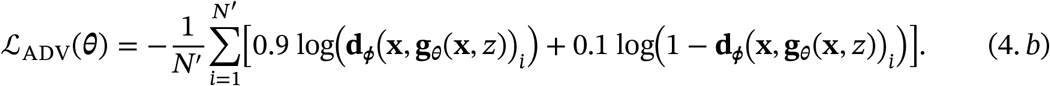

A relatively small weight λ of 2 × 10^−4^ is assigned to the adversarial loss to induce only subtle structural and textural refinements, while leaving weighted MSE still dominating the training. The generator is optimized by an Adam optimizer with a learning rate 5 × 10^−5^, *β*_1_ = 0.5, *β*_2_ = 0.999 to achieve a fine-tuning effect and maintain training stability. The final and exponential moving average (EMA) checkpoints from both training stages were retained^42^. Owing to their greater stability, the EMA checkpoints were used for evaluation.

### Input layer dropout strategy

During each training epoch, for any given 3-channel SEEQPI input, there is a probability *p* that one of its two adjacent images (either the upper or lower slice) is replaced by an all-zero image. The selection between the upper and lower slice is equiprobable. The modified input, which now contains one entirely zeroed channel, is then directly fed into the network. This forces the network to learn how to make accurate predictions even when relying on partial information, which is particularly crucial when the training dataset is small.

### Definition of evaluation metrics

Given the ground-truth and predicted binary masks, pixels were classified as true positives (TP), true negatives (TN), false positives (FP), or false negatives (FN). The following metrics are defined:

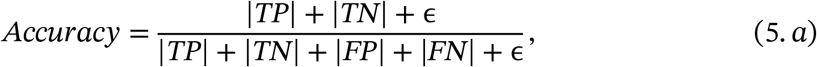

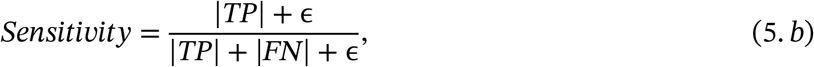

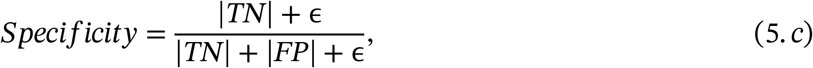

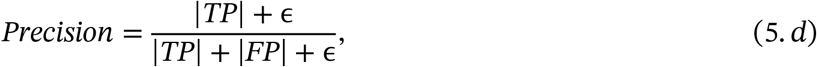

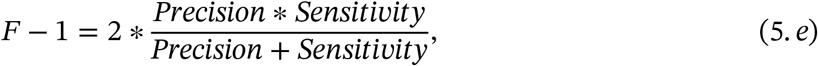

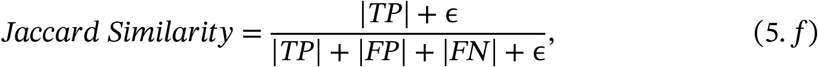

where |·| indicates the number of pixels in each category. The final and exponential moving average (EMA) checkpoints from both training stages were retained. The EMA checkpoints yielded more stable predictions.

### Dry mass density estimation

In approximation, the dry mass density *M*(**r**) has a linear relationship with the SEEQPI phase map *S*(**r**) with the following equation

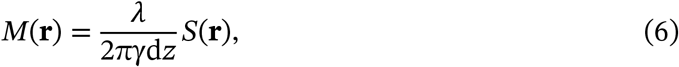

where **r** is the three-dimensional coordinate in the deep tissue^43^. λ is the central wavelength of excitation light. γ is the refractive increment, estimated as 0.2 mL g^−144^. d*z* is the imaging step size along the axial direction.

The predicted binary mask *I*(**r**) first identified the nuclear regions, within which dry-mass density was calculated using Eq. (6). Hence, the nuclear dry-mass density 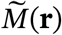 is given by:

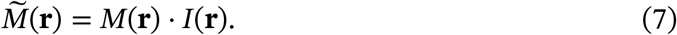

## Data availability

Due to size considerations, the data that support the findings of this study are available from the corresponding author on reasonable request.

## Code availability

The code that supports the findings of this study is available from the corresponding author on reasonable request.

## Acknowledgements

We acknowledge the support of the National Institute of Biomedical Imaging and Bioengineering (grant nos. R00EB034164 and K99EB034164 to Xi Chen).

## Contributions

X.C. proposed the idea and conceived the project. X.C. designed and built the system and performed imaging. J.Y. developed the computational algorithms, trained and evaluated the networks, and analyzed the data. J.L., N.S., and X.C. supervised network training. X.C. and J.Y. wrote the manuscript. All authors reviewed the manuscript. X.C. supervised the project.

