## Supplementary_Material for "Intelligent SEEQPI enables reconstruction of inhibitory neuronal nuclei and dry-mass mapping deep in the living mouse brain"

#### Training process visualization

Here, we present the model-training processes for the three schemes, including stage-one FiLM U-Net and stage-two FiLM U-Net + GAN. For stage one, we plot the training and validation MSE losses and sensitivities across epochs. For stage two, we show the generator and discriminator loss trajectories together with the corresponding training and validation sensitivities.

In Scheme 1, we pooled images from four mice and randomly divided them into training, validation, and test sets using three configurations: 70%/10%/20%, 60%/10%/30%, and 50%/10%/40%. For all three configurations, we performed stage-one supervised regression for 100 epochs, followed by 40 epochs of stage-two GAN training. During the first five epochs of stage two, only the generator was trained for warm-up. Figure S1–S3 show the training curves for the three configurations, respectively.

During stage one, the training weighted MSE decreases markedly, accompanied by a reduction in validation weighted MSE, suggesting that the learned mapping generalizes to unseen data. The concurrent increase in sensitivity on both sets further supports this interpretation. During stage two, both generator and discriminator losses decrease, indicating stable adversarial optimization. The generator loss declines consistently across all three configurations. For the discriminator, the 70%/10%/20% configuration shows an initial decrease followed by a plateau and a subsequent decline, whereas the other two configurations converge earlier and remain nearly stable. Nevertheless, sensitivity remains largely unchanged on both the training and validation sets throughout stage two. These results suggest that GAN fine-tuning primarily improves prediction structure and precision, with little additional gain in sensitivity.

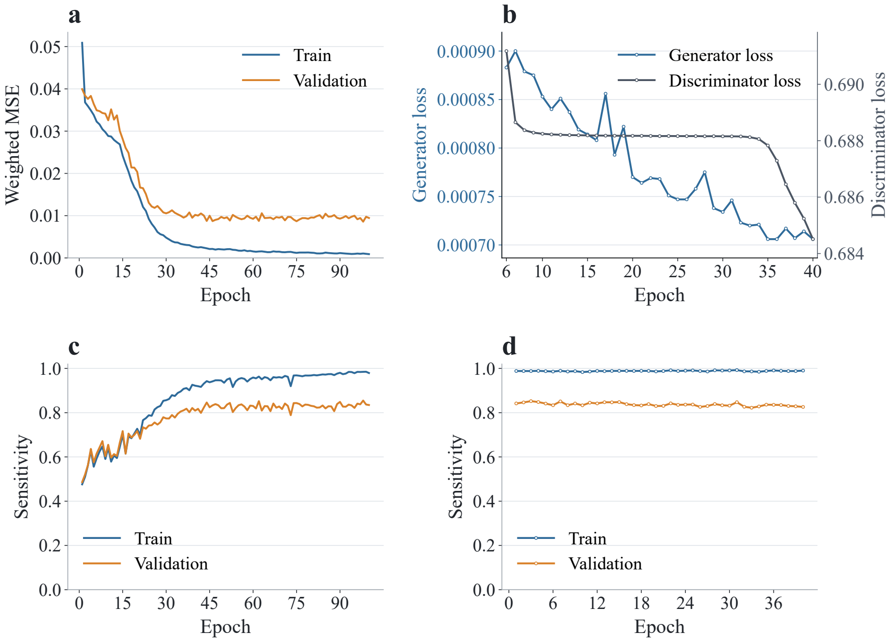


**Fig. S1. Training curves for the 70%/10%/20% training/validation/test split in Scheme 1. a,** Training and validation MSE losses during stage one. **b,** Generator and discriminator losses during stage two. **c,d,** Training and validation sensitivities during stage one (**c**) and stage two (**d**).

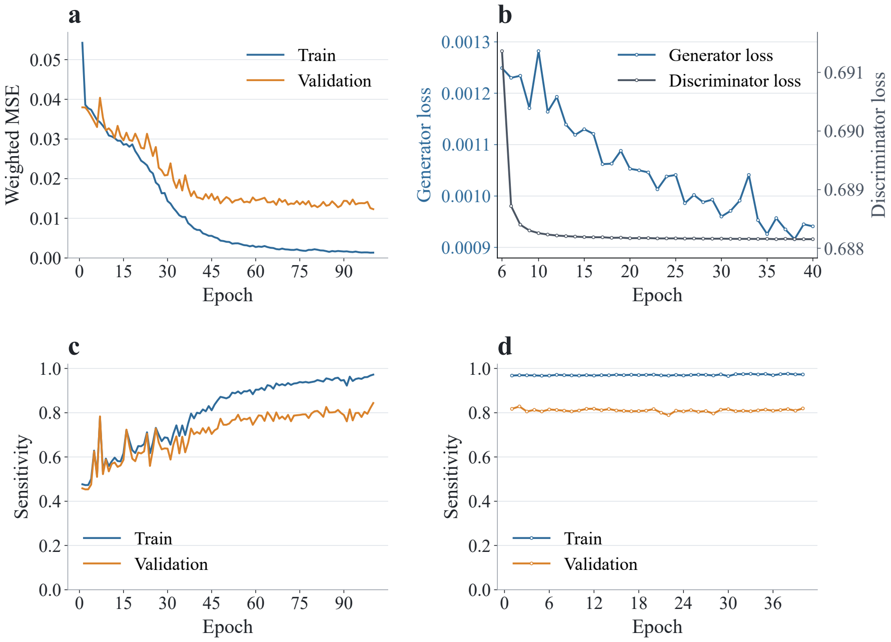


**Fig. S2. Training curves for the 60%/10%/30% training/validation/test split in Scheme 1. a,** Training and validation MSE losses during stage one. **b,** Generator and discriminator losses during stage two. **c,d,** Training and validation sensitivities during stage one (**c**) and stage two (**d**).

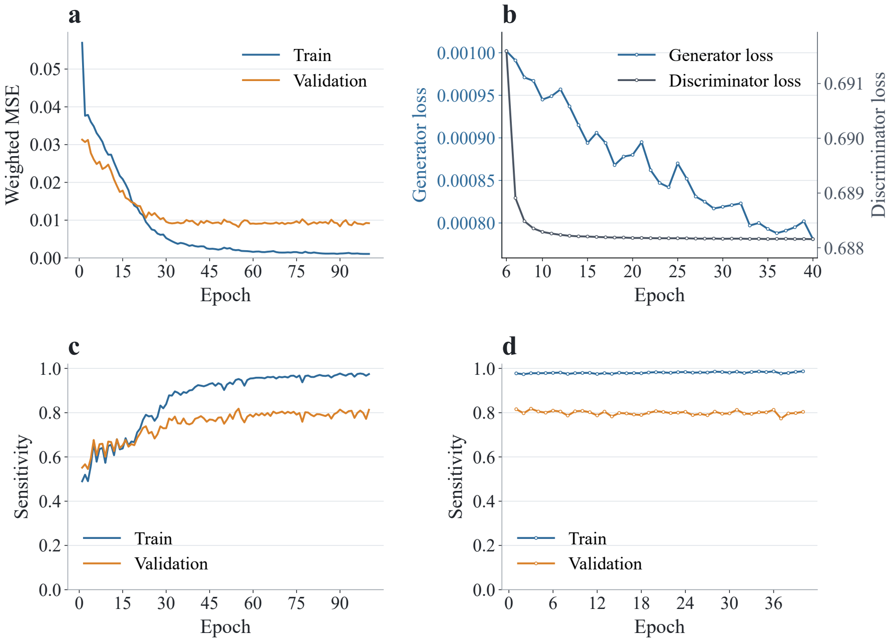


**Fig. S3. Training curves for the 50%/10%/40% training/validation/test split in Scheme 1. a,** Training and validation MSE losses during stage one. **b,** Generator and discriminator losses during stage two. **c,d,** Training and validation sensitivities during stage one (**c**) and stage two (**d**).

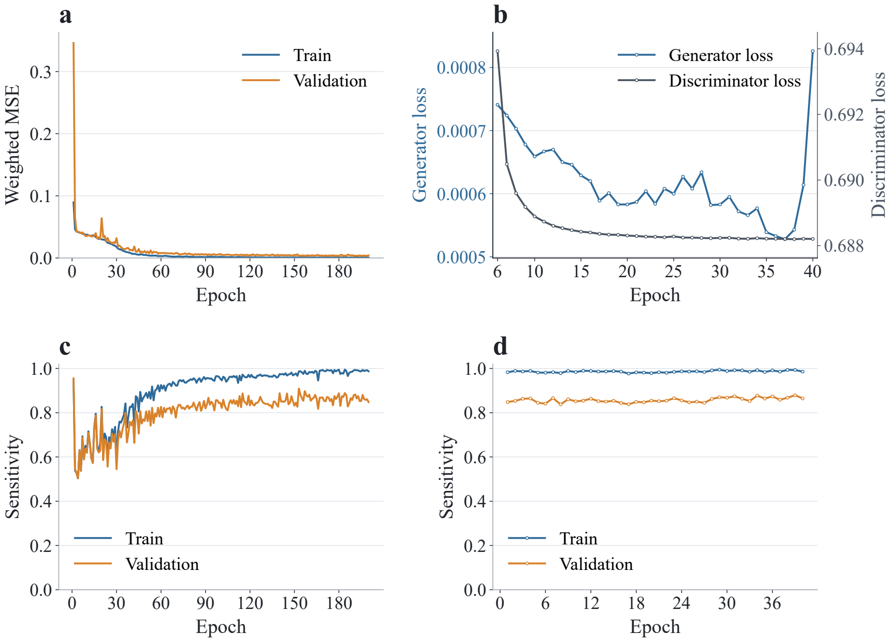


**Fig. S4. Training curves for $\Delta z = 4$ μm in Scheme 2. a,** Training and validation MSE losses during stage one. **b,** Generator and discriminator losses during stage two. **c,d,** Training and validation sensitivities during stage one (**c**) and stage two (**d**).

For the single-mouse sparse-supervision models in Scheme 2, stages one and two were trained for 200 and 40 epochs, respectively. The final EMA models were recorded. The first five epochs of stage two served as a generator-only warm-up period. Figure S4 shows the training curves at $\Delta z = 4$ µm; similar trends were observed at $\Delta z = 6 - 8$ µm. During stage one, the loss decreased markedly on both the training and validation sets. Training sensitivity increased steadily toward 1, whereas validation sensitivity initially declined before recovering. The initial decline may reflect a transition from excessive positive predictions to more conservative detection, while the subsequent increase indicates improved recovery of true nuclei. Together with the decreasing validation loss, this trend suggests that the learned features generalized to the validation data. During stage two, the discriminator loss decreased smoothly toward convergence. The generator loss also declined initially but exhibited an abrupt increase near the end of training, possibly reflecting transient adversarial instability. Nevertheless, sensitivity remained stable on both sets, suggesting that this fluctuation had little effect on nuclear detection performance.

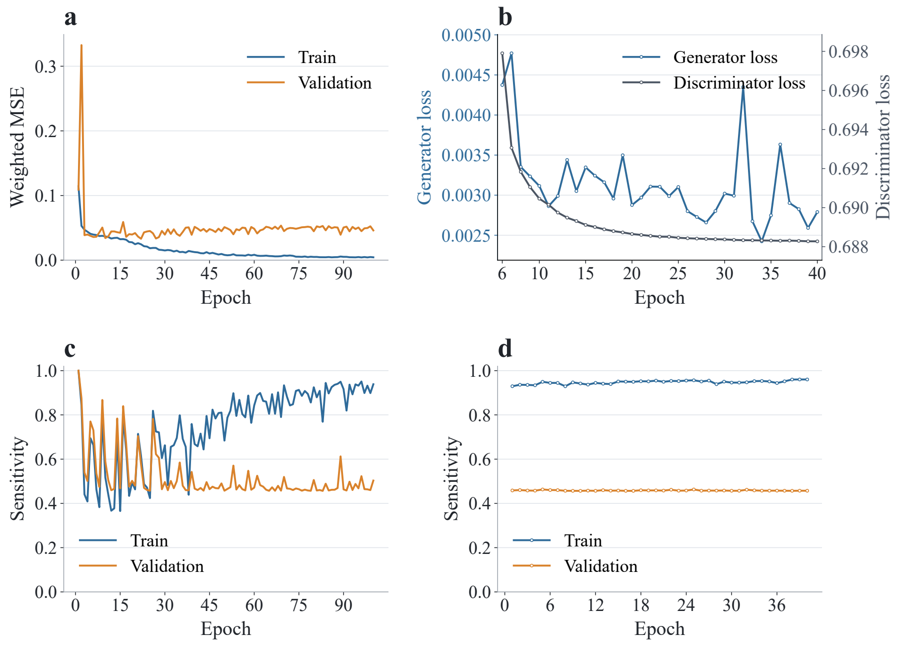


**Fig. S5. Training curves for single-stack learning on Mouse 3 in Scheme 3. a,** Training and validation MSE losses during stage one. **b,** Generator and discriminator losses during stage two. **c,d,** Training and validation sensitivities during stage one (**c**) and stage two (**d**).

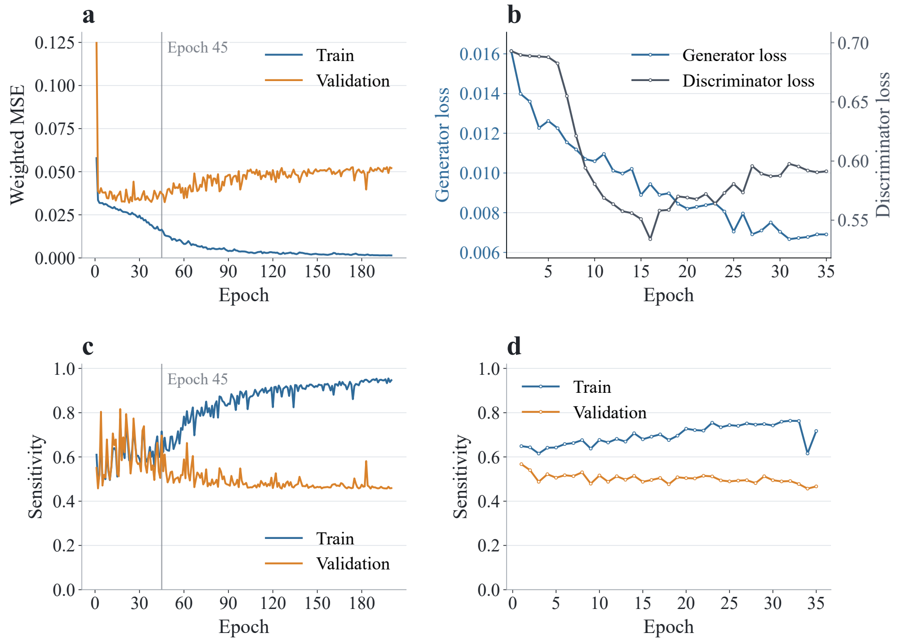


**Fig. S6. Training curves for multiple-stacks learning on Mouse 3 in Scheme 3. a,** Training and validation MSE losses during stage one. **b,** Generator and discriminator losses during stage two. **c,d,** Training and validation sensitivities during stage one (**c**) and stage two (**d**). Epoch 45 in stage one is highlighted.

In Scheme 3, Mouse 3 was selected as the prediction target, with training images sampled at an interval of $\Delta z = 8$ μm. As described in the main text, we compared the performance of models trained by single-stack and multiple-stack learning. For single-stack learning, stages one and two were trained for 200 and 40 epochs, respectively, with the first five epochs of stage two used for generator warm-up. For multiple-stack learning, early stopping was applied in both stages, and the stage-two warm-up was omitted. Figure S4 and S5 show the training curves for these two learning strategies. During stage one, the validation loss initially decreased but subsequently began to rise. To avoid both premature stopping and overfitting, we selected the checkpoint at epoch 45. During stage two, validation sensitivity declined after the first epoch; accordingly, the EMA checkpoint at epoch 1 was selected. Compared with single-stack learning, multiple-stack learning substantially improved validation-set sensitivity.

### Comparison with different neural network models

In this work, the feature-wise linear modulation U-Net (FiLM U-Net) was used for prediction in both stage-one supervised regression and as the generator in stage-two GAN training. Efficient FiLM U-Net and Attention FiLM U-Net were also evaluated as comparison models by incorporating FiLM blocks into Efficient U-Net and Attention U-Net, respectively[40,45,46]. The Efficient U-Net encoder was initialized with ImageNet-pretrained weights[47]. Following Scheme 1, slices pooled from the z-stacks of four mice were randomly divided into training, validation, and test sets at a ratio of 70%/10%/20%. For each architecture, the test-set performance of the exponential-moving-average (EMA) models obtained after each training stage is summarized in Table S1. When using the FiLM U-Net architecture, the models obtained after Stage 1 and 2 are called FiLM U-Net and FiLM U-Net + GAN in the main text.

**Table S1.** Prediction performance metrics of the FiLM U-Net, Efficient FiLM U-Net, and Attention FiLM U-Net.

| Model | Stage | Sensitivity | Precision | F1-score | Jaccard Similarity | Accuracy | Specificity |
|---|---|---|---|---|---|---|---|
| FiLM U-Net | 1 | 0.8443 | 0.7362 | 0.7502 | 0.6901 | 0.9984 | 0.9988 |
| | 2 | 0.8235 | 0.7758 | 0.7552 | 0.6993 | 0.9987 | 0.9992 |
| Efficient FiLM U-Net | 1 | 0.8287 | 0.7213 | 0.7248 | 0.6660 | 0.9982 | 0.9987 |
| | 2 | 0.7939 | 0.7304 | 0.7222 | 0.6712 | 0.9985 | 0.9991 |
| Attention FiLM U-Net | 1 | 0.8535 | 0.7102 | 0.7339 | 0.6739 | 0.9982 | 0.9986 |
| | 2 | 0.8298 | 0.7743 | 0.7674 | 0.7124 | 0.9987 | 0.9992 |

Compared with the FiLM U-Net, the Efficient FiLM U-Net yielded lower sensitivity and precision after both supervised regression and GAN fine-tuning, resulting in correspondingly lower F1 scores and Jaccard similarity. The FiLM U-Net also achieved higher accuracy and specificity.

Under supervised regression, the Attention FiLM U-Net achieved higher sensitivity but lower precision than the FiLM U-Net, while the F1 score, Jaccard similarity, accuracy, and specificity slightly favored the FiLM U-Net. After GAN fine-tuning, the Attention FiLM U-Net retained higher sensitivity and lower precision but outperformed the FiLM U-Net in F1 score and Jaccard similarity. Overall, neither architecture showed a clear performance advantage across all metrics. Given the greater complexity of the Attention FiLM U-Net, we selected the FiLM U-Net as the more appropriate architecture.

### Comparison with different training loss functions

As shown below, the training loss critically affects model performance. We compared the weighted MSE used in this study with standard MSE, a combined binary cross-entropy (BCE)–Dice loss, and its weighted variant[48].

The combined BCE-Dice loss $\mathcal{L}_{\mathrm{C}}$ is defined as

$$\mathcal{L}_{\mathrm{C}} = \alpha \cdot \mathcal{L}_{\mathrm{BCE}} + (1 - \alpha) \cdot \mathcal{L}_{\mathrm{Dice}}, \quad \text{(S1)}$$

where

$$\mathcal{L}_{\text{BCE}} = -\frac{1}{N}\sum_{i=1}^{N}[y_i \log(p_i) + (1 - y_i)\log(1 - p_i)]\,, \qquad (S2.a)$$

$$\mathcal{L}_{\text{Dice}} = 1 - \frac{2\sum_{i=1}^{N} y_i p_i + \epsilon}{\sum_{i=1}^{N} y_i + \sum_{i=1}^{N} p_i + \epsilon}. \qquad (S2.b)$$

$\alpha$, the coefficient controlling the weights between the two functions, is chosen as 0.25. $y_i$ denotes the ground-truth binary label of pixel $i$, and $p_i$ denotes the network-predicted probability that the pixel belongs to a nuclear region. $\epsilon = 10^{-7}$ is a small constant added to prevent division by zero.

To place greater emphasis on nuclear-region information during training, we proposed a weighted variant of this combined loss $\mathcal{L}_{\text{WC}}$, defined as

$$\mathcal{L}_{\text{WC}} = \alpha \cdot \mathcal{L}_{\text{W-BCE}} + (1 - \alpha) \cdot \mathcal{L}_{\text{W-Dice}}, \qquad (S3)$$

where

$$\mathcal{L}_{\text{W-BCE}} = -\frac{1}{N}\sum_{i=1}^{N} w_i \cdot [y_i \log(p_i) + (1 - y_i)\log(1 - p_i)], \qquad (S4.a)$$

$$\mathcal{L}_{\text{W-Dice}} = 1 - \frac{2\sum_{i=1}^{N} w_i y_i p_i + \epsilon}{\sum_{i=1}^{N} w_i y_i + \sum_{i=1}^{N} w_i p_i + \epsilon}. \qquad (S4.b)$$

$w_i$ is the weight assigned to each pixel. Its definition is identical to that used for weighted MSE: pixels labeled as nuclei in the ground-truth binary mask are assigned a weight of 101, whereas background pixels are assigned a weight of 1.

We again used the experimental configuration in which images from four mice were pooled and randomly split into training, validation, and test sets at a ratio of 70%/10%/20% to evaluate models trained with the four loss functions. The resulting performance metrics are presented in Table S2.

Models trained with standard MSE and the combined BCE–Dice loss exhibited the same behavior: high precision but low sensitivity, indicating that only a small proportion of nuclear signals were detected. This likely resulted from the sparse nuclear signals in the three-photon training images, which biased learning toward the dominant background and produced conservative nuclear predictions.

Compared with the unweighted BCE–Dice loss, its weighted variant substantially improved sensitivity but markedly reduced precision. The weighting encouraged the model to predict more nuclear regions but also produced numerous false-positive signals. Overall, the weighted MSE loss achieved the best balance between sensitivity and precision, yielding higher F1 scores and Jaccard similarity and the best overall performance among the four loss functions. Accordingly, it was used for model training in all subsequent experiments.

**Table S2.** Performance metrics of models trained with different loss functions.

| Loss | Model | Sensitivity | Precision | F1-score | Jaccard Similarity | Accuracy | Specificity |
|---|---|---|---|---|---|---|---|
| Weighted MSE | FiLM U-Net | 0.8443 | 0.7362 | 0.7502 | 0.6901 | 0.9984 | 0.9988 |
| | FiLM U-Net + GAN | 0.8235 | 0.7758 | 0.7552 | 0.6993 | 0.9987 | 0.9992 |
| MSE | FiLM U-Net | 0.6578 | 0.9663 | 0.6828 | 0.6501 | 0.9988 | 0.9999 |
| | FiLM U-Net + GAN | 0.7088 | 0.9611 | 0.7399 | 0.6996 | 0.9990 | 0.9999 |
| BCE-Dice | FiLM U-Net | 0.6233 | 0.8744 | 0.6385 | 0.6008 | 0.9984 | 0.9999 |
| | FiLM U-Net + GAN | 0.6314 | 0.8262 | 0.6171 | 0.5795 | 0.9984 | 0.9999 |
| Weighted BCE-Dice | FiLM U-Net | 0.7765 | 0.5544 | 0.5412 | 0.4919 | 0.9959 | 0.9966 |
| | FiLM U-Net + GAN | 0.7619 | 0.6871 | 0.6620 | 0.6130 | 0.9977 | 0.9986 |

### Effect of input-layer dropout on model performance

Input-layer dropout, as a data-augmentation strategy, was widely used during model training to improve generalization to the test set. For each three-channel SEEQPI input, either the upper or lower adjacent-channel image was replaced with an empty image with probability $p$. Here, we evaluated the effect of different dropout probabilities $p$ on model performance.

Using the pooled four-mouse dataset with a 70%/10%/20% training/validation/test split, we compared model performance at $p = 0.0$ and $p = 0.05$, while keeping all other experimental conditions unchanged. Table S3 presents the test-set performance metrics for FiLM U-Net and FiLM U-Net + GAN under the two probabilities. The model trained with input-layer dropout performed better across all test-set metrics, with particularly pronounced improvements in sensitivity and precision.

Under the single-mouse sparse supervision setting of Scheme 2, the dropout probability $p$ that yields optimal predictive performance increased with the axial spacing $\Delta z$ between adjacent training images. For each $\Delta z$ of 4, 5, 6, and 7 μm, we evaluated dropout probabilities $p$ of 0.0, 0.05, 0.10, 0.12, and 0.15, yielding 20 experimental configurations. Additionally, when $\Delta z$ is 8 μm, we tested the performance of models trained under $p$ of 0.12 and 0.15. Sensitivity, our primary metric of interest, was used to evaluate model performance under each condition. The test-set prediction results of the EMA model obtained from FiLM U-Net + GAN are summarized in Table S4.

**Table S3.** Comparison of models trained with and without input-layer dropout under the pooled multi-mouse setting.

| Dropout Probability | Model | Sensitivity | Precision | F1-score | Jaccard Similarity | Accuracy | Specificity |
|---|---|---|---|---|---|---|---|
| 0.0 | FiLM U-Net | 0.7156 | 0.7251 | 0.6664 | 0.6248 | 0.9983 | 0.9993 |
| | FiLM U-Net + GAN | 0.6933 | 0.7835 | 0.6772 | 0.6394 | 0.9985 | 0.9997 |
| 0.05 | FiLM U-Net | 0.8443 | 0.7362 | 0.7502 | 0.6901 | 0.9984 | 0.9988 |
| | FiLM U-Net + GAN | 0.8235 | 0.7758 | 0.7552 | 0.6993 | 0.9987 | 0.9992 |

**Table S4.** Effect of input-layer dropout probability $p$ on the test-set sensitivity of the EMA models from FiLM U-Net + GAN at different $\Delta z$ values. The highest sensitivity at each $\Delta z$ is shown in bold.

| $\Delta z$ \ $p$ | 0.00 | 0.05 | 0.10 | 0.12 | 0.15 |
|---|---|---|---|---|---|
| 4 | 0.8192 | **0.8620** | 0.5663 | 0.6549 | 0.7050 |
| 5 | 0.4984 | 0.7101 | **0.8017** | 0.6041 | 0.5648 |
| 6 | 0.4923 | 0.5067 | 0.5889 | **0.6453** | 0.5105 |
| 7 | 0.4926 | 0.4946 | 0.4967 | **0.5138** | 0.5113 |
| 8 | | | | **0.5151** | 0.5013 |

At every $\Delta z$, models trained with input-layer dropout achieved higher sensitivity than the corresponding model trained without dropout ($p = 0.0$). These results demonstrate a clear benefit of input-layer dropout under sparse supervision. The optimal $p$ values at $\Delta z$ of 4, 5, 6, and 7 μm were 0.05, 0.10, 0.12, and 0.12, respectively, showing an increasing trend. However, the sensitivity gains from increasing $p$ became less pronounced at larger $\Delta z$. At $\Delta z = 8$ μm, the sensitivities at $p$ of 0.12 and 0.15 were 0.5151 and 0.5013, respectively, indicating that further increasing $p$ did not improve model performance. This may be because increasing $\Delta z$ reduces the number of training images and lowers the achievable prediction performance, thereby limiting the benefit of the additional training diversity introduced by dropout.

Overall, input-layer dropout was widely adopted as a data-augmentation strategy in this work and substantially improved model generalization under both pooled multi-mouse training and single-mouse sparse supervision. During training, the upper or lower adjacent SEEQPI image was randomly replaced with an all-zero image, increasing the effective diversity of the training inputs. This stochastic perturbation regularized parameter optimization, reduced overfitting, and prevented excessive reliance on adjacent slices. The central SEEQPI image remained unchanged to preserve essential information from the target plane. Because the SEEQPI inputs were z-score normalized, the all-zero images remained statistically compatible with the other normalized input images. However, the benefit of dropout diminished with increasing $\Delta z$ in Scheme 2, indicating that it could not compensate for the reduced amount of real training data and that accurate prediction still depended on learning meaningful features from genuine SEEQPI images.

### Effect of the mask-binarization threshold on sensitivity and precision

All previously reported metrics were calculated using a fixed threshold to convert the predicted three-photon images into binary nuclear masks. Here, we further examine the effects of varying this threshold from 0 to 1 using precision–recall curves, where recall is equivalent to sensitivity.

Figure S7 presents the precision–recall curves of the EMA models trained under the three random train/validation/test split configurations in Scheme 1 (70%/10%/20%, 60%/10%/30%, 50%/10%/40%). The curves of FiLM U-Net and FiLM U-Net + GAN models are shown together for comparison. Across all configurations, the curves extend toward the upper-right corner and yield high areas under the precision–recall curve (AUPRC), substantially exceeding the random-prediction baseline defined by the positive-pixel prevalence. The AUC decreases as the training-set proportion is reduced. Compared with supervised regression, GAN fine-tuning generally favors higher precision at the expense of sensitivity, consistent with our previous observations.

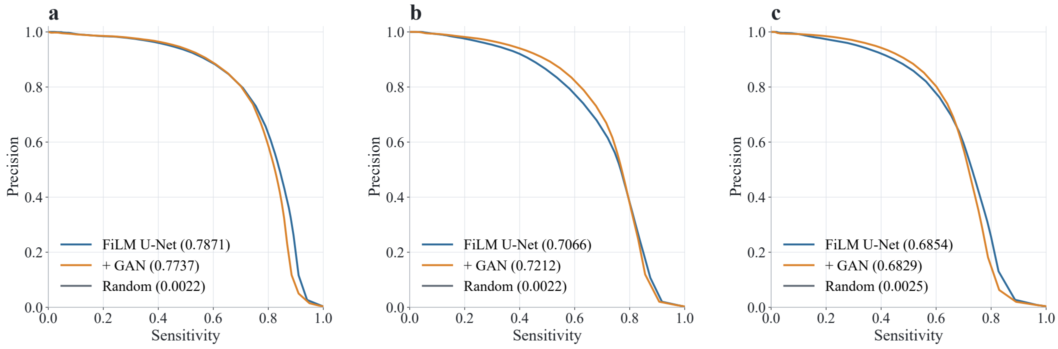


**Fig. S7. Precision–recall curves of the models trained in Scheme 1. a,** 70%/10%/20% train/validation/test split. **b,** 60%/10%/30% split. **c,** 50%/10%/40% split. Values in brackets indicate the AUPRC. + GAN represents the model FiLM U-Net + GAN.

Figure S8 shows the precision–recall curves of models trained at different $\Delta z$ values under the single-mouse sparse-supervision setting of Scheme 2. The AUPRC decreases with increasing $\Delta z$ but remains well above the random-prediction baseline, supporting the effectiveness of the method. At each $\Delta z$, FiLM U-Net + GAN achieves a higher AUPRC than FiLM U-Net, indicating that GAN fine-tuning improves the overall precision–recall trade-off.

Figure S9 compares the AUPRC values of models trained using single- and multiple-stack learning in Scheme 3. Relative to single-stack learning, multiple-stack learning increased the AUPRC by 74% for the FiLM U-Net and by 169% for FiLM U-Net + GAN, demonstrating the benefit of incorporating training data from multiple mice. Moreover, all trained models achieved AUPRC values above the random-prediction baseline.

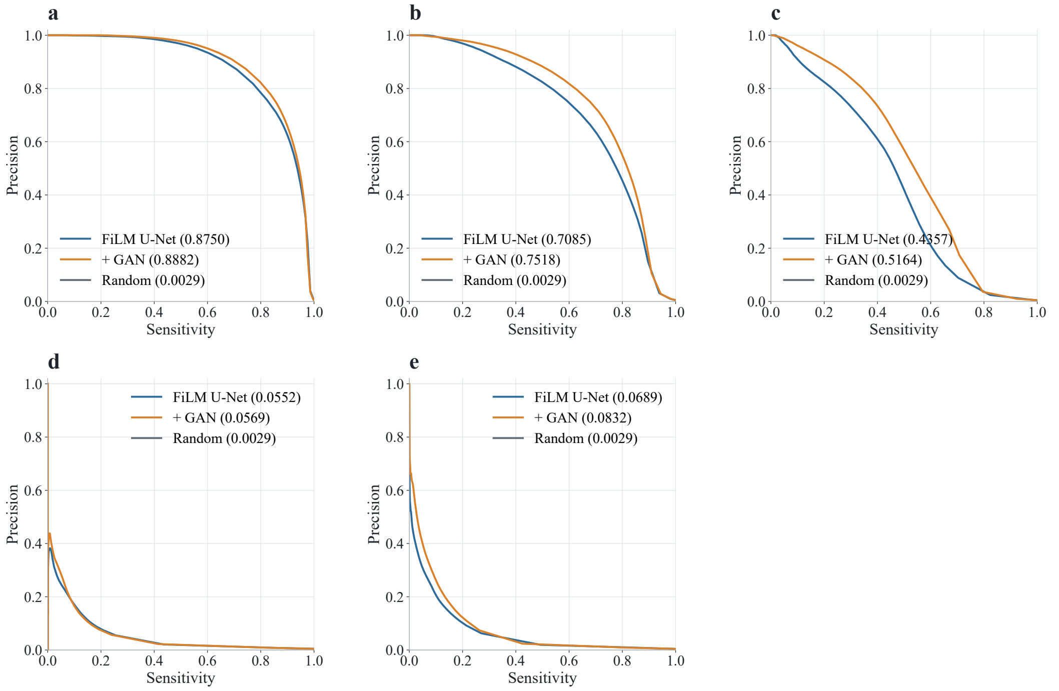


**Fig. S8. Precision–recall curves of the models trained in Scheme 2. a,** $\Delta z = 4$ µm. **b,** $\Delta z = 5$ µm. **c,** $\Delta z = 6$ µm. **d,** $\Delta z = 7$ µm. **e,** $\Delta z = 8$ µm. Values in brackets indicate the AUPRC. + GAN represents the model FiLM U-Net + GAN.

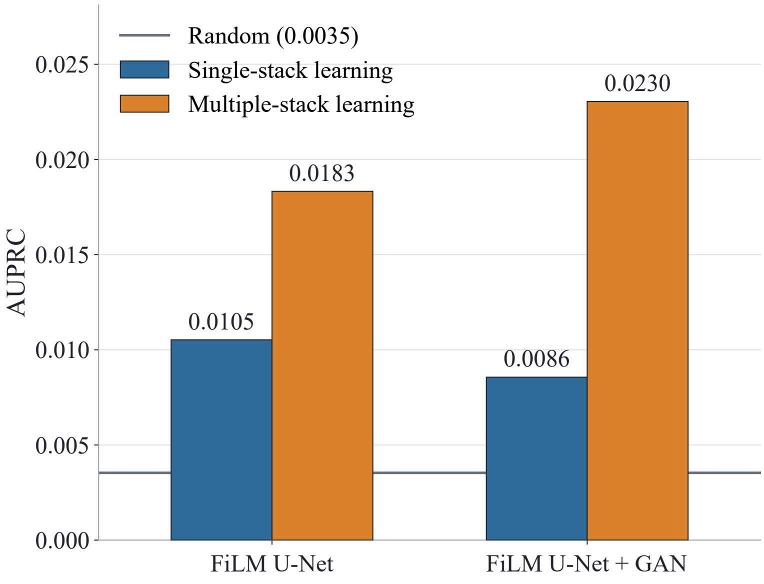


**Fig. S9.** Comparison of AUPRC on Mouse 3 between models trained using single- and multiple-stack learning in Scheme 3. Both FiLM U-Net and FiLM U-Net + GAN are included. Values in brackets indicate the AUPRC.
